# A glycan atlas of the mammalian intestine through ontogeny and inflammation

**DOI:** 10.1101/2025.03.06.641959

**Authors:** Steven J. Siegel, Daniel Pomerantz, Jamie Heimburg-Molinaro, Shaikh Danish Mahmood, Joshua R. Korzenik, Richard D. Cummings, Seth Rakoff-Nahoum

**Affiliations:** Division of Infectious Diseases, Department of Pediatrics, Boston Children’s Hospital, Boston, MA, United States; Department of Surgery, Beth Israel Deaconess Medical Center, Harvard Medical School, Boston, MA, United States; Division of Gastroenterology, Hepatology and Endoscopy, Brigham and Women’s Hospital, Boston, MA, United States; Division of Gastroenterology, Department of Pediatrics, Boston Children’s Hospital, Boston, MA, United States; Department of Microbiology, Harvard Medical School, Boston, MA, United States; Broad Institute of Harvard and MIT, Boston, MA, United States

## Abstract

The muco-epithelial interface in the mammalian gut is composed of a mucus and epithelial lining fundamental to barrier function, microbe-host interactions, and intestinal homeostasis. This barrier is heavily glycosylated by O-linked sugars covalently linked to mucin glycoproteins, and N-linked sugars that coat epithelial surface proteins. Gut O- and N-glycans are thought to play central roles in barrier function, host defense, nutrition and attachment for commensals and pathogens, immunoregulation and cell-cell interactions. However, the precise nature of the glycans and how glycan composition changes through development, as a function of diet, and during inflammation, remains incompletely understood. Here, we apply O- and N-glycomic platforms to profile glycans on mucus and intestinal epithelium. By mapping individual glycan species spatially and temporally we identify 57 O- and 18 N-glycans in the mouse intestine, and observe that fucosylation and sialylation varies according to intestinal region and developmental stage. We identify a subset of glycans regulated by the gut microbiome, and observe a constriction of the glycan repertoire during inflammation in both mice and humans. Together, these results provide an atlas of individual intestinal glycans and their dynamic range through ontogeny and inflammation, and represent a significant resource for our understanding of the role of intestinal glycans in health and disease and glycan-focused therapies for intestinal inflammation and shaping the gut microbiome.

**Highlights:**

- Individual glycans vary across gut region and developmental stage
- Terminal fucose and sialic acid residues vary across space and time
- The microbiome influences gut glycan composition early in life
- Gut inflammation in mice and humans converge on a restricted glycan repertoire

**eTOC blurb:** Microbes colonizing the mammalian intestines encounter mucus and an epithelial layer highly decorated by glycans. Siegel et al. use glycomics to map these sugars in high resolution across gut region, microbial colonization, development and inflammation in both humans and mice.

## Introduction

The intestinal surface is coated in mucus layers that protect the underlying epithelium and deeper host tissues from damage and infection while permitting physiological host processes such as nutrient absorption.^1–3^ As part of these dual functions, mucus serves as a key interface between the epithelium and the commensal microbiota. The secreted mucins that form this layer are heavily modified with glycans, chains of covalently linked carbohydrates on proteins and lipids that contribute to the gel properties of mucus.^4,5^ Glycoconjugates are assembled by combining individual monosaccharides through different attachments to the underlying protein or lipid, linkages within glycans, and with distinct extensions and terminal modifications. Each level of organization suggests the potential for significant diversity in intestinal glycans and contributes to the vast chemical diversity that glycans encode, but the precise structure and nature of the glycans in the mammalian gut have remained elusive.^6,7^

Two key glycan modifications are the addition of sialic acid and fucose to mucin glycoproteins. In the terminal position facing the intestinal lumen, these monosaccharides are the first to interact with colonizing microbes. Sialylation of the major mucin glycoprotein, MUC2, alters mucus stability and influences microbiota composition.^8^ Loss of Cosmc, a chaperone needed for function of the T-synthase glycotransferase, leads to the truncation of most mucin O-glycans, followed by significant changes in the gut microbiota and spontaneous colitis.^9^ Which glycans, among the many altered with *Cosmc* deletion, shape the microbiome remains unknown. The total amount of sialic acid and fucose varies along the intestines, and during inflammation, and host glycosyltransferases change in expression through ontogeny.^3,10–17^ These terminally-modified glycans have the potential to select for individual members of the microbiota, such as by providing specific energy sources and binding motifs for mucus-associated bacteria, but mechanistic studies at this interface have been limited by the few studies defining individual glycans in the intestinal glycome.^18^ Most studies of intestinal glycosylation have relied on genetic knockouts that impact a wide range of glycans, staining with lectins to identify classes of glycans or mass spectrometry that can identify individual glycans but only applied to purified glycoproteins. Microbial enzymes necessary for glycan foraging and binding are specific to individual glycans and linkages, making it imperative to understand the intestinal glycome in greater detail than in most existing studies.

Intestinal inflammation can be initiated by chemical or physical injury, autoimmunity and infection.^19^ Diverse inflammatory stimuli alter host glycans, such as by increasing terminal fucosylation and changing glycan extension.^11,13,14^ Inflammatory bowel disease, encompassing Crohn’s disease and ulcerative colitis, is an idiopathic disease of intestinal inflammation that involves a loss of mucosal barrier function.^20–22^ Glycosylation is key to this barrier, but individual glycans present in the secreted mucus of patients with IBD remain unknown.^23–26^ Here, we apply glycomics to chart an atlas of mucus and epithelial O- and N-glycans in mice and humans.

## Results

### Intestinal glycans are region-specific

Previous studies of the intestinal glycome have focused on a single species, a limited number of tissue sites, or both. To overcome these limitations and evaluate different compartments within the intestine, we modified O- and N- glycomics platforms for application to limited mass samples such as mucus and epithelial tissue. We first surveyed the small and large intestine of adult B6 mice reared in specific-pathogen free conditions (**Figure 1A**). After removing luminal contents (stool), we used mechanical disruption to harvest the adherent inner mucus layer, followed by chemical treatment with EDTA to isolate the underlying epithelium. From each intestinal compartment, we extracted O- and N-glycans from glycoproteins using chemical (alkaline β-elimination) and enzymatic (PNGase F) methods, respectively. These glycans were permethylated and their structures identified using MALDI-TOF mass spectrometry. Using these methods, here we report an atlas of intestinal glycans across development, species, gut region and disease (**Figure 1B**).

**Figure 1.**
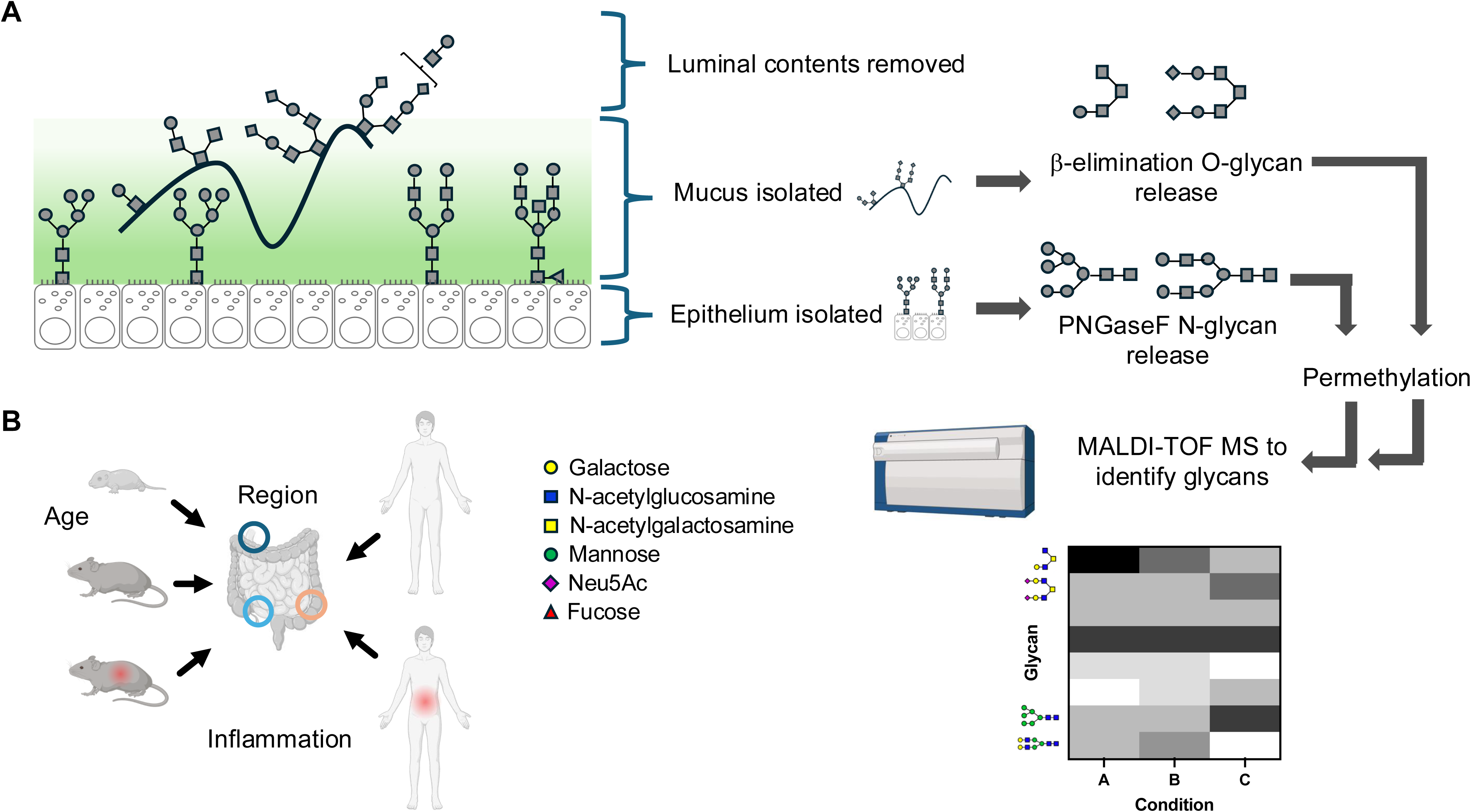
Pipeline for intestinal glycomics. (A) The workflow for isolating and analyzing mouse intestinal glycans is depicted. Mouse duodenum, ileum and colon were isolated at necropsy. Each intestinal segment was opened longitudinally, luminal fecal material was removed and the mucus layers mechanically separated from the underlying epithelium. The epithelial layer was isolated from the underlying mucosa using EDTA. Mucus fractions were subjected to chemical release of O-glycans, while N-glycans were enzymatically released from the epithelial fraction using PNGaseF. O- and N-glycans were purified and analyzed by MALDI-TOF mass spectrometry. (B) The scope of the glycan atlas presented here is depicted, including mouse age, gut region (duodenum in dark blue, ileum in light blue and colon in orange), species and inflammation.

We applied this pipeline to define host glycosylation along the longitudinal axis of the gut. Analyzing the spectra generated (**Figure 2A**), we identified 57 O- and 18 N-glycans in the mucus and epithelium, respectively, of wildtype adult mouse duodenum, ileum and colon, from 3-6 mice pooled per sample type (**Tables S1-2**). We categorized each O-glycan by its terminal modifications, the monosaccharide(s) most accessible to colonizing microbes in the gut lumen. The most prominent such modifications in the mammalian gut are sialic acid, fucose, the combination of both sialic acid and fucose, or neither (**Figure 2B**). These residues can also be added to glycans in nonterminal positions, underscoring the importance of glycan-specific methods of identification.^27^ Prior studies demonstrated opposing gradients of sialic acid and fucose in the murine gut, with sialylation decreasing from proximal to distal gut while fucosylation increased, though the changes in each terminal modification were not quantified. We measured the change in relative intensity of sialylated and fucosylated glycans, finding a reduction in the proportion of sialylated as well as dually non-sialylated, non-fucosylated glycans in the distal gastrointestinal tract compared to the proximal gut (**Figure 2C**). We therefore identified an increasing proportion of terminally modified O-glycans from proximal to distal gut. Separate from terminal modifications, the proximal end of O-glycans determine their categorization into one of several core subtypes. Annotating each glycan identified by MALDI-TOF, we measured the relative abundance of each core glycan subtype; the majority of O-glycosylation on the mucus layer was core 2 (**Figure 2D**), across the whole longitudinal axis of the gut.

**Figure 2.**
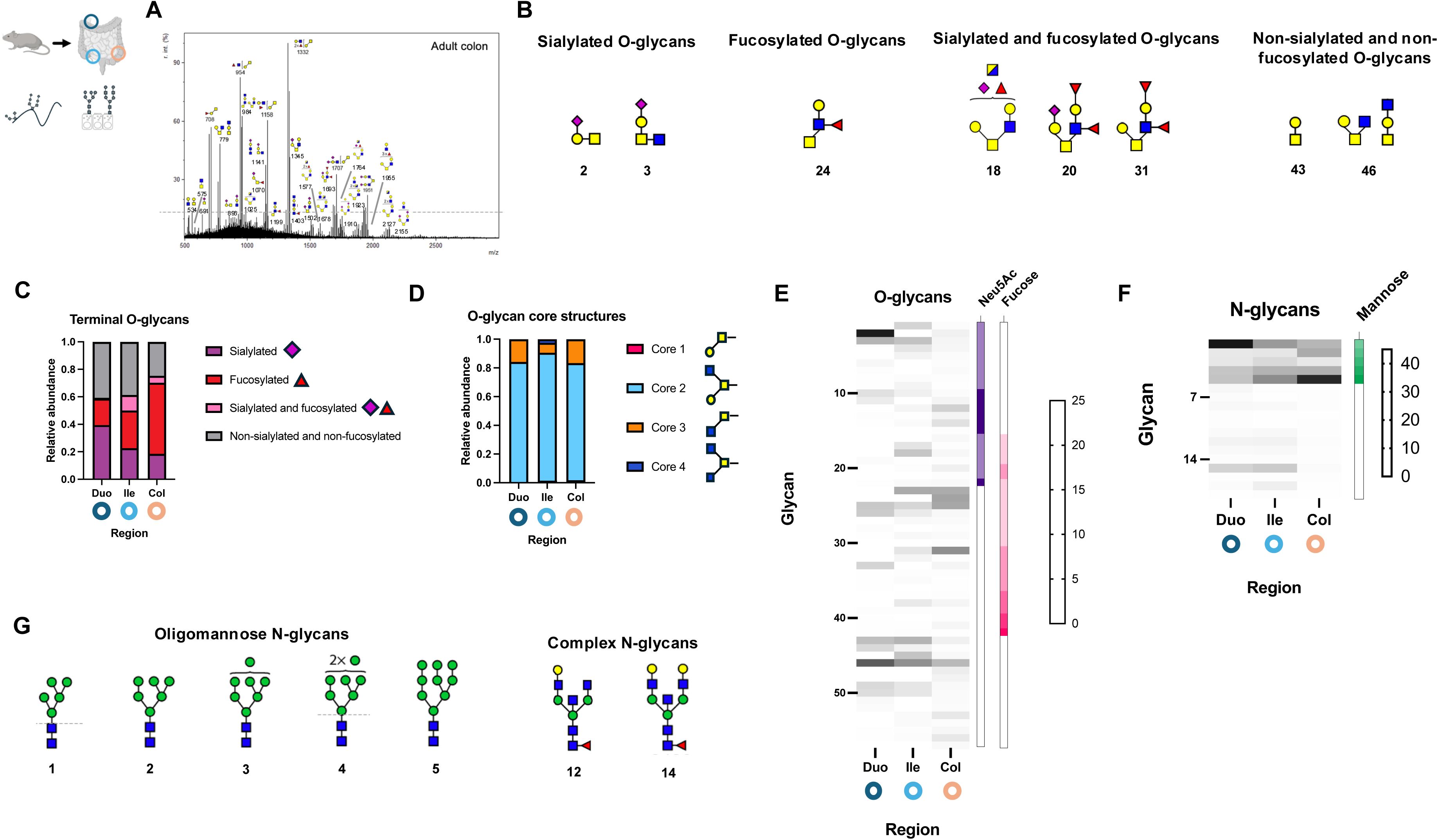
Intestinal glycans are region-specific. (A) O-glycans were isolated and purified from the mucus layers of mouse duodenum, ileum and colon, and analyzed by MALDI-TOF. A mass spectrum from one of the colon samples is presented, with annotated glycans noted by mass and predicted structures displayed. (B) Predicted glycan structures identified in the mouse colon are displayed according to category of terminal modification. Numbers below each structure refers to their order in subsequent heat maps and in **Table S1**. (C) The relative proportion of glycans in each terminal modification category is displayed according to gut region. (D) The relative proportion of glycans predicted to be extensions of each core O-glycan is displayed according to gut region. (E) The relative abundance of each identified O- glycan within a gut region is depicted in a heat map. Each row represents a different glycan identified by MALDI-TOF, and each column denotes a different intestinal region. The number of sialic acid (Neu5Ac) and fucose residues predicted in each identified glycan is depicted by the intensity of shading in the boxes to the right of the heat map. (F) The relative abundance of each identified N-glycan within a gut region is depicted in a heat map. Each row represents a different glycan identified by MALDI-TOF, and each column denotes a different intestinal region. The number of mannose residues predicted in each identified glycan is depicted by the intensity of shading in the box to the right of the heat map. (G) Predicted N-glycan structures identified in mouse epithelial samples are displayed according to category. N = 5-10 pooled mice per condition.

We next asked whether changes in overall sialylation and fucosylation reflected changes in all glycans with those terminal modifications or only a subset. To address this question, we constructed heat maps to assess changes in the abundance of each glycan at each region of the gut (**Figure 2E**). We observed distinct patterns of abundance changes. Some sialylated glycans (such as glycan 2) decreased in intensity across the gut, while others (such as glycan 12) increased, even as total sialylation decreased. Similarly, distinct abundant fucosylated glycans in the colon increased in expression compared to the duodenum while another subset decreased in abundance. Our findings therefore suggest individual glycan-level changes in terminal modifications rather than a complete shift from sialylation to fucosylation.

We next analyzed the N-glycans isolated from the gut epithelium (**Figure 2F-G**). The majority of all N-glycans were oligomannose rather than complex, suggesting that most N-glycans present were precursors rather than the more modified complex type. While the individual N-glycan species varied by region, N-glycan classes remained dominated by oligomannose rather than complex structures at all sites. These results suggest distinct mechanisms for regulating the glycans that decorate mucus and the epithelium.

### Intestinal glycans are specific to developmental stage

Limited studies of human fetal intestines suggested differences in intestinal glycosylation compared to adults. To test if glycosylation varies during development, we compared intestinal mucus O-glycan abundances from infant (10-12 day old), just-weaned (24-28 day old) and adult (42 day old) mice (**Figure 3A**). Within each intestinal region (duodenum, ileum and colon), we found substantial differences across age groups. A smaller number of O-glycans dominated the duodenum with increasing age, though without substantial changes in the relative abundance of sialylation. Within the ileum, sialylation dominated more at younger ages, with a transition to more mixed sialylation and fucosylation during development. In the colon, O-glycans that are mostly fucosylated in adult mice are instead more mixed among sialylation, fucosylation and neither terminal modification in infants (**Figure 3B**). The more diverse range of terminal glycan modifications in the immature colon was mirrored by core glycan distributions. Infant colonic mucus glycans were distributed across a wider range of core subtypes than in adults, with core 2 O-glycans showing the most abundance (**Figure 3C**).

**Figure 3.**
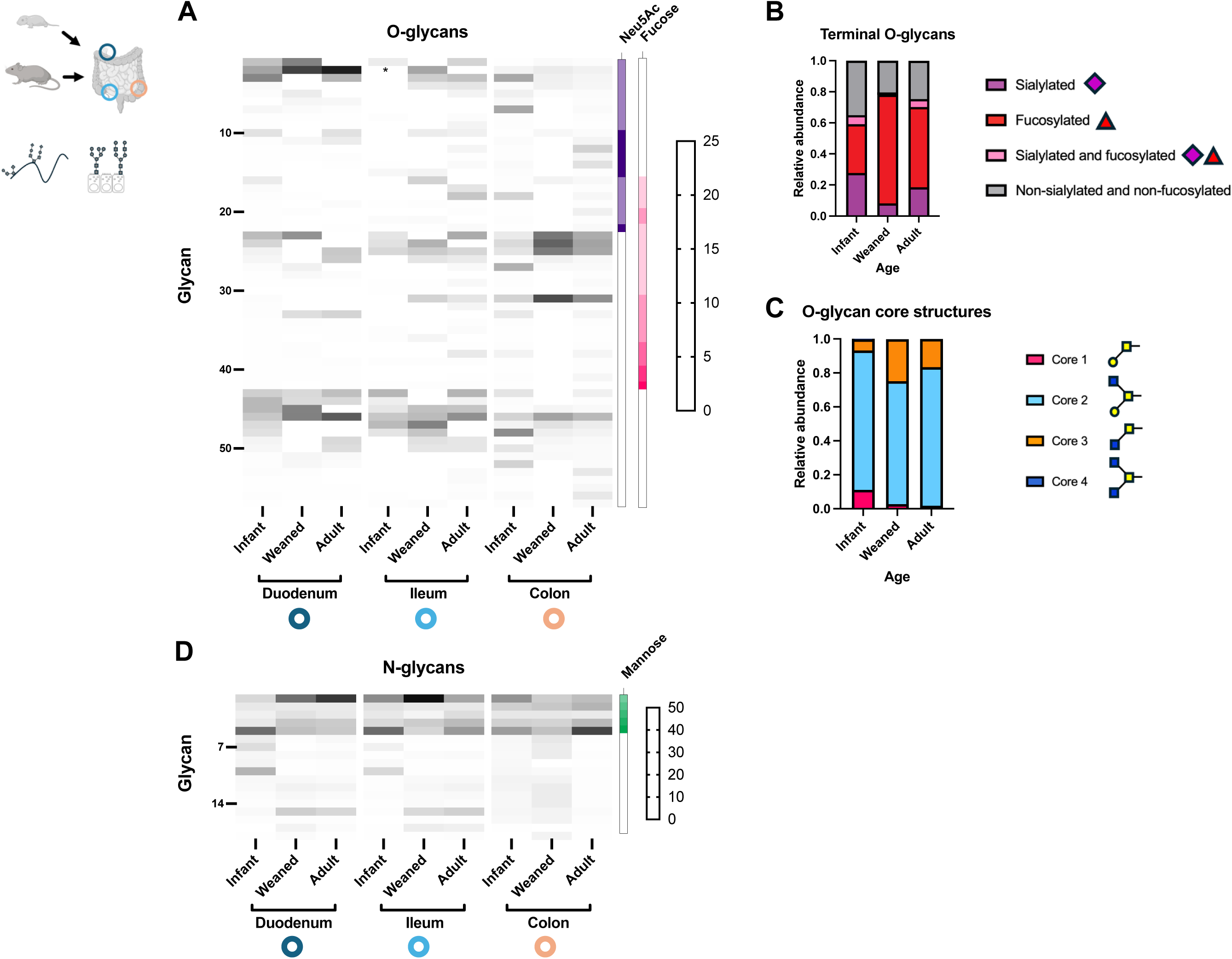
Intestinal glycans are specific to developmental stage. (A) The relative abundance of each identified O-glycan within a gut region and developmental stage is depicted in a heat map. Each row represents a different glycan identified by MALDI-TOF, and each column denotes a different intestinal region at a different age mouse. The number of sialic acid (Neu5Ac) and fucose residues predicted in each identified glycan is depicted by the intensity of shading in the boxes to the right of the heat map. The asterisk indicates intensity above the highest value indicated on the heat map legend. (B) The relative proportion of glycans in each terminal modification category is displayed according to developmental stage. (C) The relative proportion of glycans predicted to be extensions of each core O-glycan is displayed according to developmental stage. (D) The relative abundance of each identified N- glycan within a gut region and developmental stage is depicted in a heat map. Each row represents a different glycan identified by MALDI-TOF, and each column denotes a different intestinal region at a different age mouse. The number of mannose residues predicted in each identified glycan is depicted by the intensity of shading in the box to the right of the heat map. N = 3-12 pooled mice per condition.

We next asked whether N-glycosylation also varied according to developmental stage and intestinal region. In the duodenum and ileum, complex N-glycans were relatively unchanged between weaned and adult mice, but distinct from infant mice (**Figure 3D**). This pattern was not present in the colon, where N-glycans shifted by adulthood to more exclusively oligomannose-type. Importantly, however, the overall dominance of oligomannose N-glycans in the gut was unchanged regardless of developmental stage or region, unlike in mucus O-glycans.

### The microbiome shapes intestinal glycan composition

We next examined the mechanisms regulating developmental and regional changes in intestinal glycosylation. Colonic mucus O-glycans were largely unchanged after weaning from maternal milk. The process of weaning and switching diet influences the composition of the microbiome, which becomes more stable and reaches its final, adult form during adolescence.^28^ To test whether gut-resident microbes alter intestinal glycosylation, we compared glycosylation between mice fed a standard (control) or polysaccharide-free (PS-free) diet for 1 week prior to analysis. The latter limits dietary carbohydrates to simple monosaccharides, driving gut microbes towards scavenging host glycans from the mucosal surface. The relative abundance of fucosylated glycans increased slightly under a PS-free diet (e.g., decrease in glycan 14 and increase in glycan 18), but overall patterns of glycan expression and terminal glycan modifications were unchanged (**Figure 4A**). These results suggested that microbial complex glycan consumption is not a substantial driver of mucus glycan expression. To more directly test the contribution of the microbiome (whether as glycan consumers, editors or as immune stimulants) to intestinal glycosylation, we applied our glycomics pipeline to control (specific-pathogen free) and germ-free mice at each developmental stage and across each intestinal region. In the duodenum of infant mice, multiple O-glycans decreased in relative frequency in the absence of a commensal microbiome, while glycan 2 was substantially increased (**Figure 4B**). Polyfucosylated glycans increased in abundance in the duodenum in germ-free weaned and adult mice, suggesting these glycans were under different microbial regulation. In the ileum of infants, by contrast, a wider range of glycans were expressed in the absence of the microbiome (**Figure 4C**). The most substantial differences were found in the colon of conventional compared to germ-free mice (**Figure 4D**). In the absence of a microbiome, germ-free infant mice revert to a glycosylation profile more similar to that of weaned or adult mice. We tested how similar O-glycosylation was among groups by principal component analysis, demonstrating that conventionally-raised infant mice have a unique glycome (**Figure 4E**). These results suggest that the infant-specific microbiome regulates mucus O- glycan expression.

**Figure 4.**
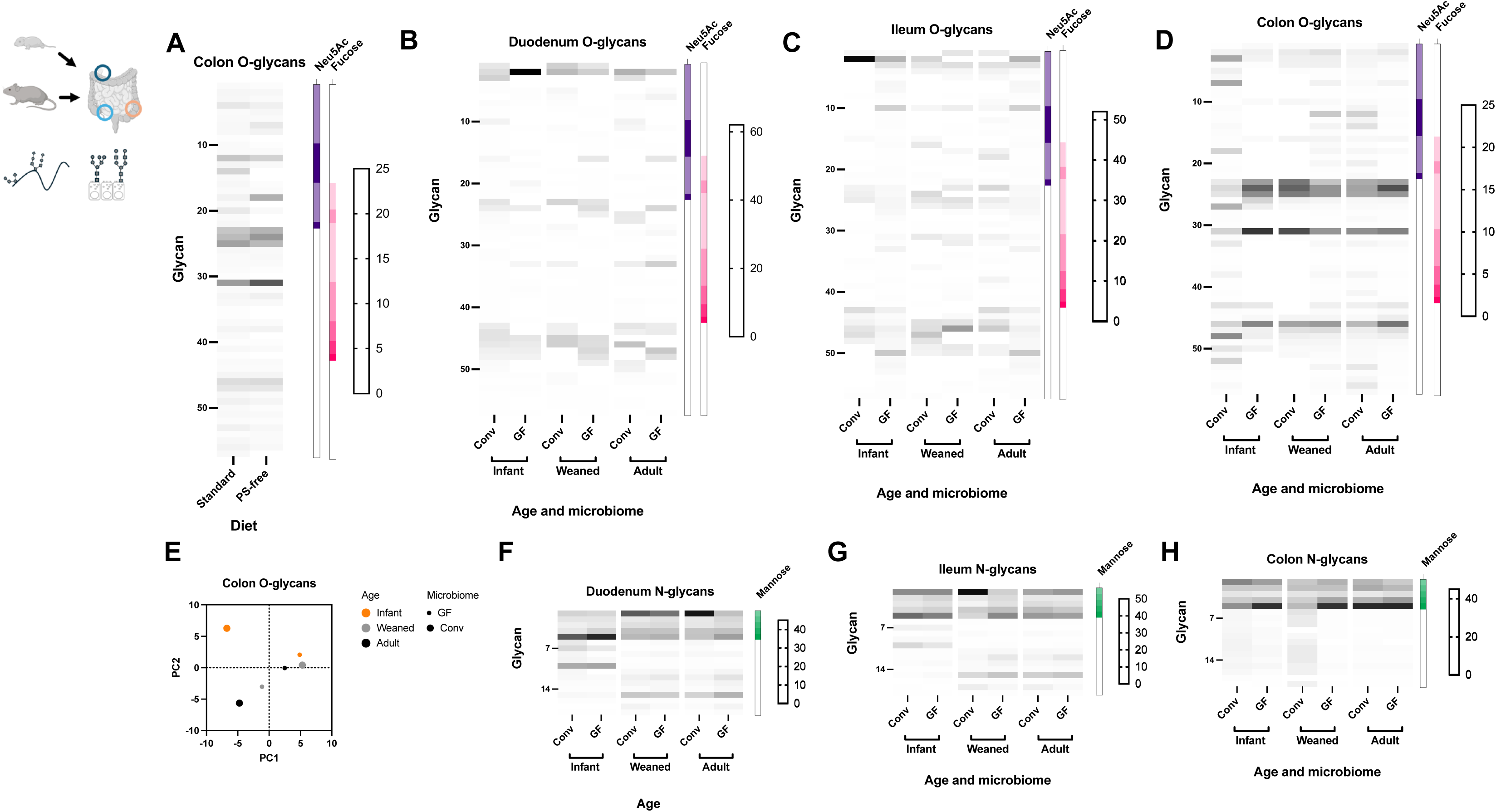
The microbiome shapes intestinal glycan composition. (A) Mice were given either standard chow or a polysaccharide-free diet for 10 days prior to glycan isolation. The relative abundance of each identified O-glycan according to mouse diet is depicted in a heat map. Each row represents a different glycan identified by MALDI-TOF, and each column denotes a different diet. Standard indicates standard mouse chow and PS-free indicates polysaccharide diet. The number of sialic acid (Neu5Ac) and fucose residues predicted in each identified glycan is depicted by the intensity of shading in the boxes to the right of the heat map. The relative abundance of each identified O-glycan according to age and microbiome status in the (B) duodenum, (C) ileum and (D) colon is depicted in heat maps. Each row represents a different glycan identified by MALDI-TOF, and each column denotes a different developmental stage and microbiome status. Conv denotes mice raised in a conventional, specific-pathogen free environment. GF denotes mice raised in a germ-free environment. The number of sialic acid (Neu5Ac) and fucose residues predicted in each identified glycan is depicted by the intensity of shading in the boxes to the right of the heat map. (E) Principal component analysis was used to display the relative similarity of mouse colon mucus O-glycan distributions according to developmental stage and microbiome status. The relative abundance of each identified N-glycan according to age and microbiome status in the (F) duodenum, (G) ileum and (H) colon is depicted in heat maps. Each row represents a different glycan identified by MALDI-TOF, and each column denotes a different developmental stage and microbiome status. Conv denotes mice raised in a conventional, specific-pathogen free environment. GF denotes mice raised in a germ-free environment. The number of mannose residues predicted in each identified glycan is depicted by the intensity of shading in the box to the right of the heat map. N = 5-12 pooled mice per condition.

We hypothesized that N-glycosylation of the intestinal epithelium would not be under microbial regulation given its distance from the luminal microbiome. In the duodenum and ileum, N-glycan abundances were largely unchanged between conventional and germ-free animals (**Figures 4F-G**). By contrast, in the colon of infant and weaned mice, germ-free animals had fewer complex N-glycans and a higher relative abundance of oligomannose glycans (**Figure 4H**). These results suggested more limited microbial regulation of epithelial N-glycans than mucus O-glycans.

### Intestinal inflammation constricts the glycan repertoire

Previous studies have described increased fucosylation during intestinal inflammation, but the specific glycans altered during inflammation have been unclear. To address this, we used a standard mouse model of Th1-mediated inflammation, chemical colitis from oral administration of dextran sodium sulfate. We compared mucus O-glycosylation between mice fed DSS or control. Consistent with prior reports, terminal fucosylation increased during inflammation (**Figure 5A**). Examining the relative abundance of each glycan, however, we found this increase in fucosylation was due to the substantial increase in abundance of a small number of glycans, notably glycans 24 and 31 (**Figure 5B**). This increase was specific to polyfucosylated rather than monofucosylated glycans (**Figure 5C**), consistent with an increase in expression of more complex glycans. The overall mucus glycome was restricted to a smaller number of O-glycans identified during inflammation, quantified by comparing the richness and evenness of glycan intensities between groups. The Shannon diversity index, a measure of the richness and evenness of species within a sample, decreased during inflammation, consistent with a restricted array of glycans (**Figure 5D**).

**Figure 5.**
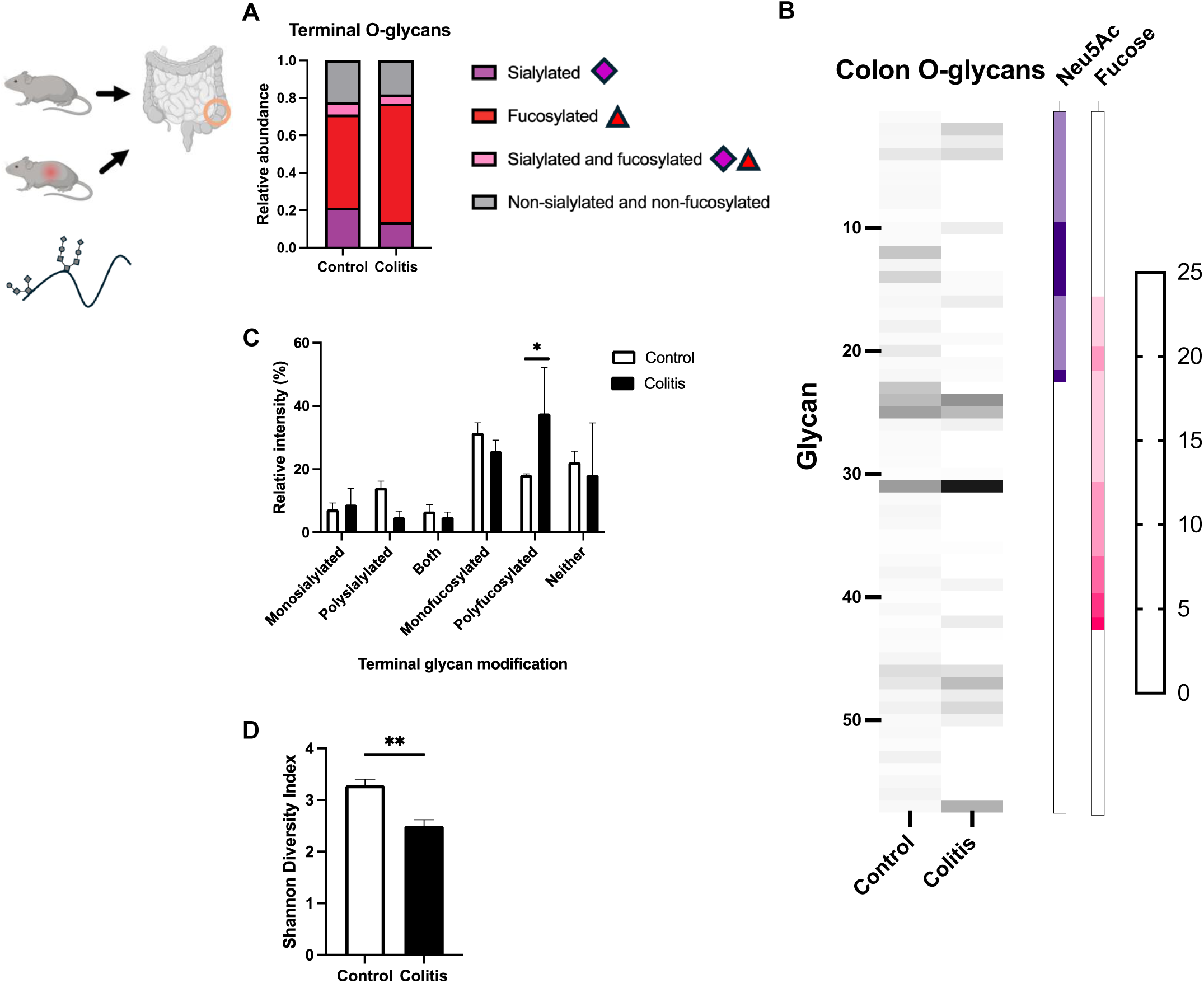
Intestinal inflammation constricts the glycan repertoire. (A) Mice were treated with 2% dextran sodium sulfate (DSS) or control for 7 days prior to glycan isolation. The relative proportion of glycans in each terminal modification category is displayed according to treatment group. (B) The relative abundance of each identified O-glycan within each treatment group is depicted in a heat map. Each row represents a different glycan identified by MALDI-TOF, and each column denotes a treatment group (colitis refers to DSS treatment). The number of sialic acid (Neu5Ac) and fucose residues predicted in each identified glycan is depicted by the intensity of shading in the boxes to the right of the heat map. (C) The relative abundance of each identified O-glycan within each treatment group is displayed according to categories of terminal modifications. Bars are mean +/- SD. *, p < 0.05. (D) The distribution of glycan abundances was compared using the Shannon diversity index. Bars are mean +/- SEM. **, p < 0.01. N = 4-8 pooled mice per condition.

### Human mucus O-glycans vary by region

Having charted an atlas of the mouse intestinal O- and N-glycome across region, age, microbiome and inflammation, we next sought to apply these techniques to the human intestinal glycome. Prior human studies have relied on intestinal biopsies,^29^ which could reflect incompletely processed mucus prior to release onto the epithelial surface. To overcome this limitation, we developed a sampling protocol using brushings of the colonic surface, followed by recovery of collected mucus for O-glycomics. We applied this protocol to healthy adults undergoing routine screening colonoscopies for colorectal cancer. Unlike mouse samples, in which 3-6 mice were pooled to generate each glycan profile, we were able to obtain sufficient material from individual brushings to profile the glycome of each subject separately. Comparing across two regions of the human gut, the ascending and descending colon, we found a total of 88 O-glycans expressed (**Figure 6A**). Some of these glycans were enriched or relatively decreased in each region examined, such as glycans 1, 26, 38, 62 and 72. These glycans were nearly equally distributed among each class of terminal modification: fucosylated, sialylated, both and neither, with no substantial gradient from ascending to descending colon (**Figure 6B**). These results suggested a more even distribution of O-glycans in human mucus compared to murine mucus that is more dominated by a single terminal modification, fucosylation.

**Figure 6.**
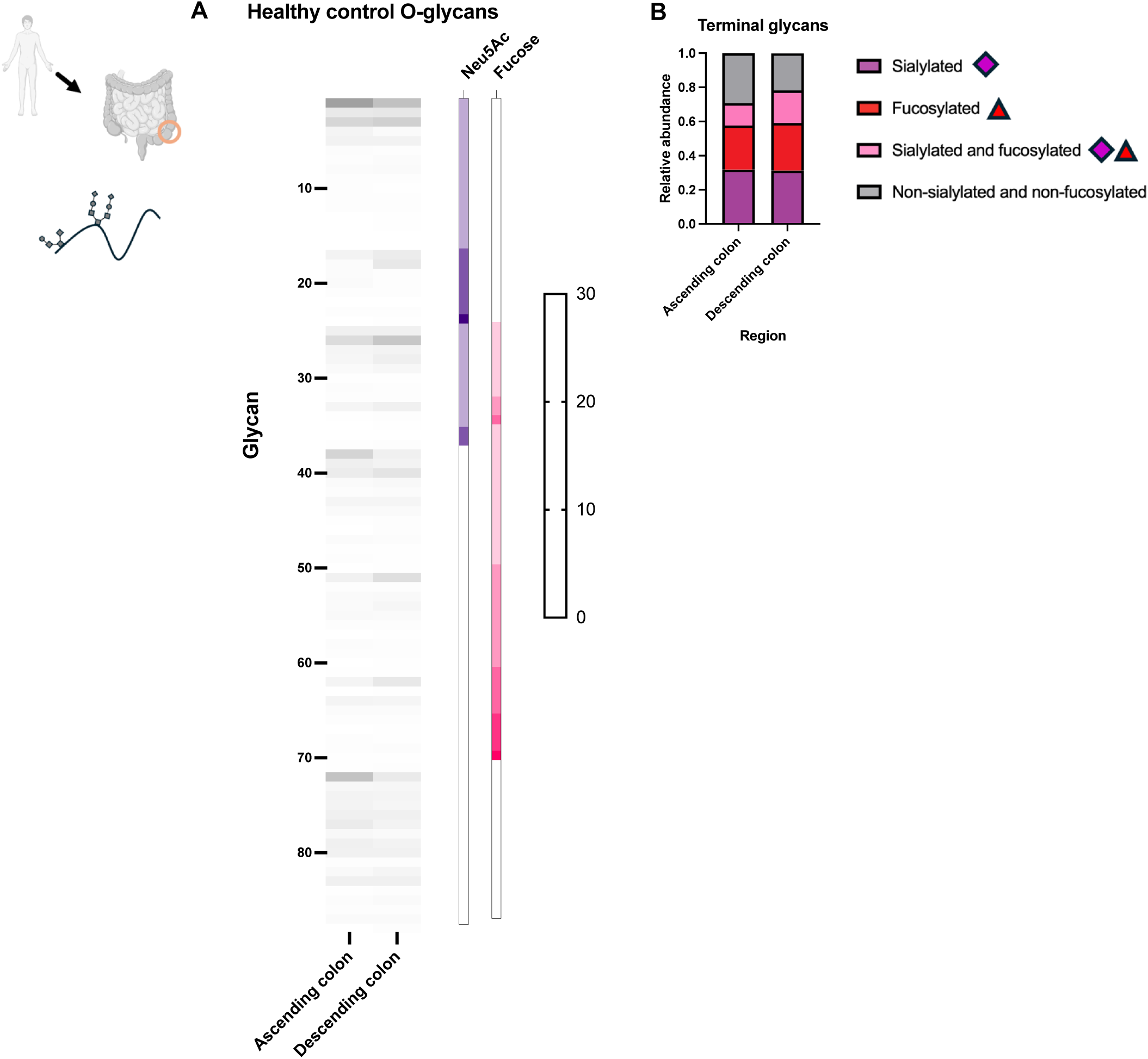
Human mucus O-glycans vary by region. (A) Mucus brushings were obtained during screening colonoscopies in healthy human patients, from the ascending and descending colon. O-glycans were isolated and identified by MALDI-TOF. The relative abundance of each identified O-glycan within each intestinal region is depicted in a heat map. Each row represents a different glycan identified and each column denotes a region. The number of sialic acid (Neu5Ac) and fucose residues predicted in each identified glycan is depicted by the intensity of shading in the boxes to the right of the heat map. (B) The relative proportion of glycans in each terminal modification category is displayed according to treatment group. N = 5 individual human samples per region.

### Glycan repertoire constriction is conserved in human inflammatory bowel disease

Prior studies have reported increased expression of smaller, less complex O-glycans in patients with inflammation, though these results stemmed from analysis of the colon epithelium itself, rather than secreted mucus.^13^ Taking advantage of our ability to obtain mucus brushings, we analyzed O-glycans in mucus isolated from patients with ulcerative colitis, a form of inflammatory bowel disease limited to the colon. We sampled 5 patients with active colitis and 5 patients without active colitis (inactive disease), each at 2 sites, the ascending and descending colon. A shift towards smaller, less complex glycans during active inflammation was observed in the descending but not the ascending colon (**Figure 7A**). We also noted a similar regional specificity when analyzing terminal modifications during active inflammation. In the ascending colon, monosialylated glycans became less abundant during active inflammation, while in the descending colon the major shift was towards an increase in non-sialylated, non-fucosylated glycans, less complex in modifications (**Figure 7B**). As in the mouse model, inflammation led to a constriction of the glycan repertoire (**Figure 7C**), demonstrating conserved features of the mucus glycan response during inflammation.

**Figure 7.**
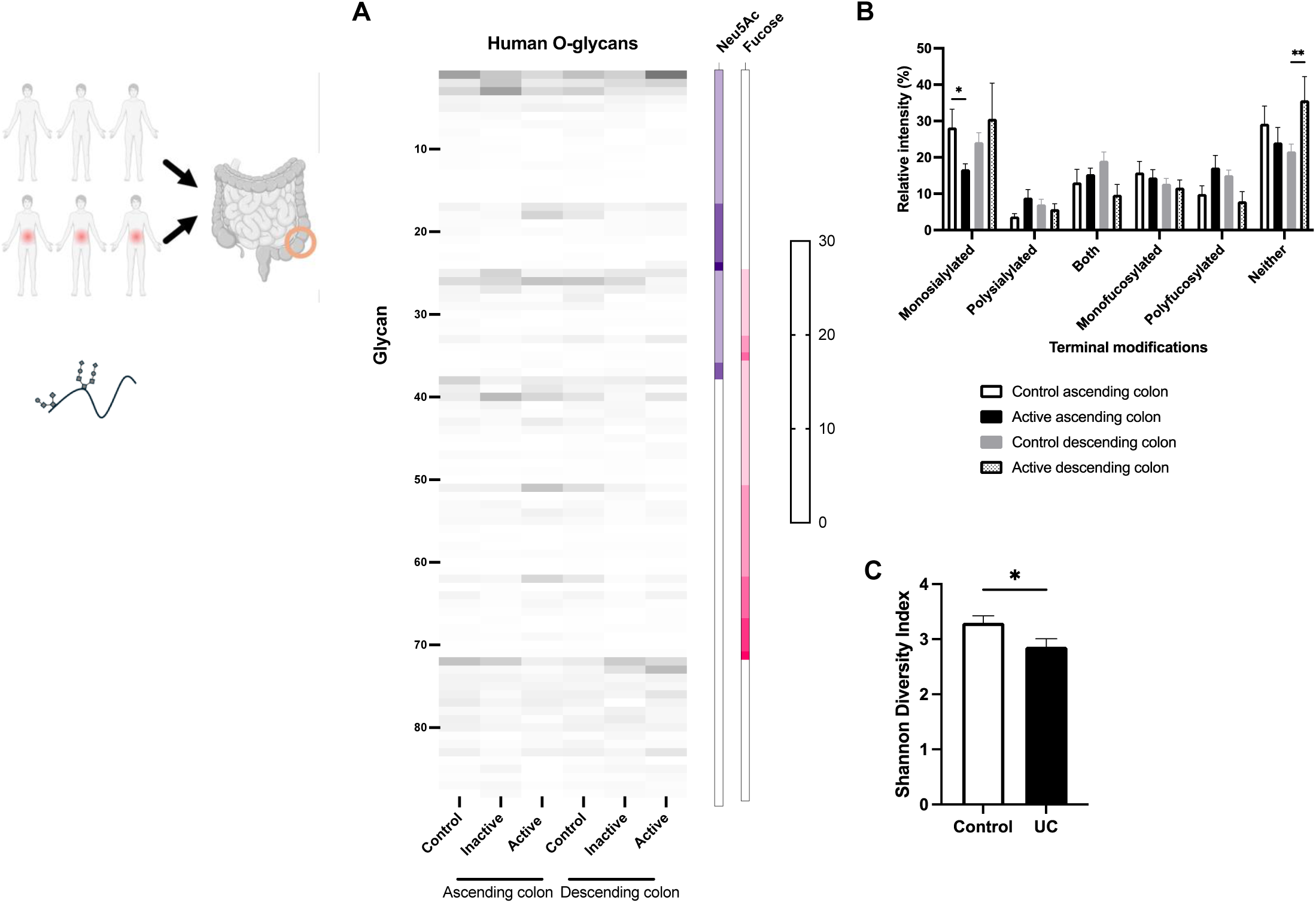
Glycan repertoire constriction is conserved in human inflammatory bowel disease. (A) Mucus brushings were obtained during screening colonoscopies in healthy individuals as well as surveillance colonoscopies in patients with active or inactive ulcerative colitis, from the ascending and descending colon. O-glycans were isolated and identified by MALDI-TOF. The relative abundance of each identified O-glycan within each intestinal region is depicted in a heat map. Each row represents a different glycan identified and each column denotes a region and patient group. The number of sialic acid (Neu5Ac) and fucose residues predicted in each identified glycan is depicted by the intensity of shading in the boxes to the right of the heat map. (B) The relative abundance of each identified O-glycan within each treatment group is displayed according to categories of terminal modifications. Bars are mean +/- SD. *, p < 0.05, **, p < 0.01. (C) The distribution of glycan abundances was compared using the Shannon diversity index. Bars are mean +/- SEM. N = 5 individual human samples per patient group.

## Discussion

Here, we present the first atlas of the intestinal mucus and epithelial O- and N-glycome in two species, across region, development, presence of the microbiome and inflammation. We identified terminal mucus O-glycan modifications under developmental and microbial regulation, revealing candidate glycans that may shape intestinal and microbiome homeostasis. By contrast, we found that epithelial N-glycans remain dominated by oligomannose rather than complex N-glycans across each of these variables. We also determined that inflammation in both mice and humans restricts the complexity of the mucus O-glycan repertoire.

It is noteworthy that the mucus O-glycome of infant mice was most distinct from that of other ages. Prior to weaning, the mammalian diet comprises milk, rich in oligosaccharides that resemble the mucin oligosaccharides of adult animals.^30–32^ Colonizing microbes consume these glycans using similar machinery, suggesting convergence between milk and mucin glycans could aid in maintenance of the microbiome. It is unclear in this model why infant mucus glycans would be distinct from that of adults. It is also unclear whether the broader distribution of O-glycans in infants could select for different microbes – aiding in microbiome assembly and development – or be due to the presence of different microbes editing, consuming and stimulating production of glycans differently.

Microbial influences could account for the difference in regulation between mucus O-glycans and epithelial N-glycans. The mucus O-glycans that face the intestinal lumen are the first to interact with colonizing bacteria, while the epithelial surface is relatively protected from direct contact. These effects may explain why N-glycans remain largely unchanged across region, development and microbiome status. The microbiome could have distal effects on glycosylation, however, as shown by a recent study identifying glycosylation changes in the brain in gnotobiotic mice colonized with human stool.^33^

Inflammation drives several glycosylation changes in mice and humans, including increased terminal fucosylation and altering glycan truncation or extension.^34,35^ Here, we demonstrate increased polyfucosylation of mucus O-glycans in the inflamed mouse colon, identifying specific glycans responsible for inflammatory fucosylation. By contrast, and similar to prior studies of colon biopsies, intestinal inflammation in humans leads to increased simpler, smaller glycans. Common to both mice and humans is a skewing of the glycan repertoire towards a smaller number of individual glycans expressed. One hypothesis for this common pathway is attempting to limit the range of glycans available to intestinal microbes that could be pathogenic during barrier disruption, a form of nutritional immunity.^36–38^

Importantly, we observe individual glycans changing in abundance independent of overall trends in terminal modifications. That is, even as fucosylation increases along the longitudinal axis of the gut, some individual sialylated glycans increase in relative abundance. These patterns underscore the importance of methods that can trace individual glycans. Some recent work has largely relied on sampling whole tissues or biopsy samples, while here we isolate mucus and epithelial cells.^39,40^

Identifying specific glycan structures that change in abundance between intestinal sites and under different conditions enables future mechanistic studies aimed at decoding the glycan-bacteria interactions that contribute to commensal bacterial binding, persistence and growth within the intestinal tract, and into how specialization for host glycans drives microbiota assembly.^41,42^ These studies could lead to a mechanistic understanding of how microbes compete and cooperate within hosts, which will lead to new, rational blueprints for selecting pre- and probiotics.^43^ Furthermore, these studies could test the theoretical concept of host control, which proposes that mammalian hosts actively control their commensal microbial communities.^44^

## Acknowledgments

This work was supported by NIH grants R24GM137763, R01GM140201 (to R.D.C.), K08AI130392, DP2GM136652, R01AI158814, R01AI171100, R01AI126915, R01DK138023 (to S.R.-N.), K12HD000850 and T32AI155391 (to S.J.S.) and T32GM007748 (to D.J.P.), the Fred Lovejoy Resident Research and Education Award, Boston Children’s Hospital Office of Faculty Development/Basic & Clinical Translational Research Executive Committees Faculty Career Development Fellowship and the Jeff Gordon Children’s Foundation PTCTC New Investigator Award (to S.J.S.), as well as the Glycoscience Fund at BIDMC and the Mass Life Science Center Research Infrastructure Grant (to R.D.C).

## Author Contributions

Experimental design (S.J.S., J.R.K., R.D.C. and S.R.-N.), performing experiments (S.J.S., D.P. and S.D.M.), data analysis and interpretation (S.J.S., D.P., R.D.C. and S.R.-N.), manuscript writing and editing (S.J.S., D.P., J.H.-M., J.R.K, R.D.C. and S.R.-N.)

## Declaration of Interests

No conflicts of interest to declare.

## Methods

### Experimental Model and Study Participant Details

#### Animals

C57BL/6 mice from Taconic were used for all experiments involving specific-pathogen free animals. SPF and germ-free mice were maintained and bred at the Brigham and Women’s Hospital Center for Comparative Medicine facility, in accordance with a protocol approved by the Brigham and Women’s Hospital IACUC. Male and female mice were used for all experiments except DSS treatment, as prior experiments demonstrated a sex-specific effect on colitis.^19^ Mice were euthanized by CO_2_ asphyxiation. Adult mice were maintained in the animal facility for at least 7 days prior to use in any experiments.

#### Human Participants

Subjects provided informed consent and were enrolled in a protocol approved by the Brigham and Women’s Hospital IRB to obtain mucus brushings during colonoscopies planned for clinical indications: colorectal cancer screening for healthy patients and surveillance or disease staging for patients with ulcerative colitis. There were 3 male and 2 female healthy control participants, 1 male and 4 female participants with active UC, and 1 male and 4 female participants with inactive UC. Patients were enrolled as a convenience sample of patients presenting to Brigham and Women’s Hospital for colonoscopy.

### Method Details

#### Mucus Scraping

After euthanasia, mouse duodenum, ileum and colon were isolated individually and opened longitudinally. Visible fecal material was removed with forceps. The mucus layers were scraped from the underlying epithelium using a fresh, sterile #15 blade scalpel and dissolved in 1 mL of ultrapure water. Pooled samples were lyophilized. For human mucus brushings, a bottle brush was gentled scraped against the wall of each indicated region during clinical colonoscopies until it became saturated with mucus. Samples were frozen until further processing could occur, in which brushes were boiled in ultrapure water for 10 minutes to liberate the mucus, followed by lyophilization.

#### Epithelial Cell Isolation

After mucus scraping, intestinal segments were immersed in cold PBS and washed three times. They were incubated with rotation in 5 mM EDTA in PBS for 25 minutes at room temperature, recovered on ice for 5 minutes, then shaken and vortexed to separate epithelial cells from underlying tissue. This tissue was removed, then the epithelial cells pelleted by centrifugation before resuspending in PBS.

#### O-glycan Isolation

O-glycans were isolated according to protocols from the National Center for Functional Glycomics (available here: https://research.bidmc.org/ncfg/protocols). Briefly, lyophilized mucus samples were resuspended in 55 mg/mL NaBH_4_ in fresh 0.1 M NaOH and incubated at 45°C overnight (14-16 hours). β-elimination was terminated by adding 4-6 drops of pure acetic acid to each reaction. A small-scale desalting column was made by loading Dowex 50W X8 (mesh size 200-400) resin into a TopTip (Glygen) and washing in 5% acetic acid. Neutralized samples were loaded onto the column, washed with 5% acetic acid and eluted by centrifugation. Samples were then vacuum concentrated and lyophilized. The samples were resuspended in 1:9 acetic acid:methanol and dried under a stream of nitrogen. Co-evaporation was repeated 3 times. A 50 mg C18 Sep-Park column was conditioned with methanol, 5% acetic acid, 1-propanol and 5% acetic acid. The dried sample was resuspended in 50% methanol and loaded onto the conditioned C18 column. The columns were washed with 5% acetic acid and the flow-through collected. Samples were then vacuum concentrated and lyophilized.

#### N-glycan Isolation

N-glycans were isolated according to protocols from the National Center for Functional Glycomics (available here: https://research.bidmc.org/ncfg/protocols). Briefly, epithelial cells were pelleted, then resuspended in ice-cold lysis buffer (25 mM TRIS base, 150 mM NaCl, 5 mM EDTA and 0.5% (w/v) CHAPS, pH 7.4). The cells were sonicated with 4-5 10 second pulses using a QSonica Q700 microtip, and then dialyzed using a 1-5 kDa molecular cutoff filter against 50 mM ammonium bicarbonate at 4°C for 16h-24h, changing the buffer twice. Reactions were lyophilized and then resuspended in fresh 2 mg/mL 1,4-dithiothreitol in 0.6 M TRIS buffer pH 8.5 and incubated at 50°C for 2 hours. An equal volume of 12 mg/mL iodoacetamide in 0.6 M TRIS buffer pH 8.5 was added, and the reactions incubated at room temperature in the dark for 2 hours. Samples were again dialyzed against 50 mM ammonium bicarbonate at 4°C for 16h-24h, changing the buffer 3 times. The samples were then lyophilized again, then resuspended in 500 µg/ml TPCK-treated trypsin in 50 mM ammonium bicarbonate and incubated overnight at 37°C. Reactions were halted with 2 drops of 5% acetic acid. A 500 mg C18 Sep-Pak column was conditioned with methanol, 5% acetic acid, 1-propanol and 5% acetic acid. Each trypsin-digested samples were loaded onto a C18 column, the columns washed in 5% acetic acid and the glycopeptides eluted with successive washes with 20%, 50% and 100% 1-propanol. Eluted fractions were vacuum concentrated and lyophilized.

Samples were then resuspended in 50 mM ammonium bicarbonate and PNGaseF added. Reactions were incubated at 37°C for 4 hours, then additional PNGaseF was added for an overnight incubation at 37°C. Reactions were terminated with 2 drops of 5% acetic acid. A 50 mg C18 Sep-Pak column was conditioned with methanol, 5% acetic acid, 1-propanol and 5% acetic acid and the PNGaseF-digested samples were loaded onto the column. The column was washed with 5% acetic acid, and the flow-through vacuum concentrated and then lyophilized for later permethylation.

#### Glycan Permethylation

Purified O- and N-glycans were permethylated according to protocols from the National Center for Functional Glycomics (available here: https://research.bidmc.org/ncfg/protocols). Briefly, a fresh slurry of NaOH/DMSO was made in a mortar and pestle, and this slurry was added to lyophilized samples. Iodomethane was added and the sample was incubated at 4°C for 2 hours followed by 30 minutes at room temperature. The reaction was stopped with 5% acetic acid, and chloroform was added. The samples were vortexed, then centrifuged briefly to separate aqueous and chloroform phases. The aqueous layer was discarded and the chloroform fraction washed 3 times with water. The chloroform fraction was collected, vacuum-concentrated and dissolved in 50% methanol, then loaded onto a 50 mg C18 Sep-Pak column conditioned with methanol, water, acetonitrile and water. The column was washed with water and 10% acetonitrile, then eluted with 50% acetonitrile, followed by vacuum concentration and lyophilization.

#### MALDI-TOF Mass Spectrometry

Permethylated glycans were dissolved in 75% methanol and 10 mg/mL 2,5 DHB, then spotted on an MTP 384 polished steel target plate. FlexControl software was used to acquire spectra using a Bruker Ultraflex II mass spectrometer. Twenty independent captures, each with 1000 shots, were obtained for each sample. Spectra were annotated using mMass software.^45^

#### Quantification and Statistical Analysis

Details of the number of samples tested are in each figure legend. Graphpad Prism (10.2.3) was used for statistical analysis. Mann-Whitney U test and 2-way ANOVA were used to compare the medians of different conditions for single and multiple group comparisons, respectively. Precision measures (SD and SEM) and how significance was defined are detailed in each figure legend.

## Notes

### Competing Interest Statement

The authors have declared no competing interest.

### Summary of Updates

This version of the manuscript has been revised to update the funding information.

